# The AltR transcription factor responds to plant thiosulfinates to regulate gene expression in a bacterial pathogen of onion

**DOI:** 10.1101/2025.11.19.689302

**Authors:** Hsiao-Hsuan Jan, Feng Kong, Michelle P. MacLellan, Li Yang, Bhabesh Dutta, Brian H. Kvitko

**Affiliations:** University of Georgia, Department of Plant Pathology, Carleton St., Athens, GA 30602, USA; University of Georgia, Department of Plant Pathology, Rainwater Rd., Tifton, GA 31793, USA

**Keywords:** AltR, TetR-family repressor, thiosulfinates, allicin, *Pantoea ananatis*, onion center rot, bacterial stress response, thiol toxicity, host-pathogen interactions

## Abstract

*Pantoea ananatis,* the causative agent of onion center rot, encounters potent antimicrobial thiosulfinates, volatile organosulfur compounds released from damaged *Allium* tissues. The allicin tolerance (*alt*) gene cluster allows *P. ananatis* to overcome this chemical barrier. We demonstrate that AltR, a TetR-family transcriptional repressor, specifically regulates expression of the *alt* cluster and thus thiosulfinate tolerance *in vitro* and fitness *in vivo*. We identified a putative AltR binding box both in the *altR* promoter and elsewhere in the *alt* cluster, show that AltR-mediated repression is relieved in response to thiosulfinates. Using cysteine to serine substitutions, we demonstrate that AltR Cys100 is essential for thiosulfinate-responsive de-repression, while other AltR cysteine residues tune responsivity. Strains expressing AltR alleles with reduced thiosulfinate responsivity have reduced fitness *in planta*. Our findings uncover a regulatory mechanism by which a plant antimicrobial secondary metabolite acts as an environmental cue to modulate bacterial gene expression, enabling pathogen survival and virulence.

## Introduction

Onions (*Allium cepa* L.) are among the most economically important vegetable crop in the United States of America, valued at approximately 1 billion (USD) annually between 2018 and 2022 ^1^. Onion center rot (OCR), caused by several *Pantoea* species including *Pantoea ananatis* in the southeastern U.S., is a major bacterial disease, particularly in the Vidalia onion-growing region of Georgia ^2,3^. OCR typically begins as water-soaked lesions on leaves that progress to blighting and tissue collapse. Bulb symptoms may remain latent at harvest and develop during storage ^4–6^. No onion center rot resistant onion cultivars are currently available, and disease management relies primarily on copper bactericides and insecticides targeting thrips which can serve as vectors ^7–10^.

Unlike many plant-pathogenic bacteria, *P. ananatis* lacks the canonical virulence-associated type II and type III secretion systems commonly required for plant infection^11–13^. Instead, *P. ananatis* strains that cause OCR produce the phosphonate phytotoxin pantaphos, which is required for necrosis and symptom development in onion ^14–17^. Although the mode of action for pantaphos remains unclear, exposure of onion tissue to pantaphos leads to cell death within days ^18^.

Upon cell death and membrane disruption, onions release thiosulfinates; thiol-reactive, sulfur-containing phytoavengins ^19–21^. In intact cells, non-toxic S-alk(en)yl cysteine sulfoxides (CSOs) accumulate in the cytoplasm, while alliinase (CSO lyase) resides in the vacuole. Upon tissue disruption, CSOs mix with alliinase, producing sulfenic acid intermediates that spontaneously condense into thiosulfinates. These volatile compounds contribute to characteristic aroma of *Allium* species and exhibit strong antimicrobial activity by forming mixed-disulfides with cellular thiols including proteins cysteine residues which can inactivate critical enzymes ^22–29^.

Successful *P. ananatis* OCR strains encode the *alt* (allicin tolerance) thiosulfinate tolerance gene (TTG) cluster, or which enables colonization of necrotic onion tissue ^30^. The *alt* cluster confers tolerance to both endogenous onion thiosulfinates and the garlic-delivered thiosulfinate allicin ^31^. Homologous TTG clusters have been identified in other onion-associated *Pantoea* species, and similar clusters have been identified several onion-pathogenic *Burkholderia* species and in the garlic saprophyte *Pseudomonas fluorescens* PfAR-1 ^32–35^.

Among the *alt* genes, several encode enzymes are predicted to detoxify or reduce thiosulfinate stress ^30,36^. A single transcriptional regulator, AltR, is conserved across characterized a*lt* clusters and is predicted to encode a TetR-family repressor ^30,31^. TetR-family repressors are known to be self-regulating homodimeric repressors controlling genes involved in secondary metabolism, antibiotic resistance, or stress responses, including the canonical TetR that confers tetracycline resistance ^37^. These regulators feature an N-terminal helix-turn-helix (HTH) DNA-binding domain and a C terminal ligand-binding/dimerization domain ^38,39^. In the *Escherichia coli* bleach responsive repressor NemR (an AltR homolog), one conserved critical redox-active cysteine was suggested as a key residue that controls responsiveness ^40^.

In this study, we provided genetic evidence delineating the regulatory role of *P. ananatis* AltR in thiosulfinate tolerance. We show that AltR represses its own promoter through an inverted repeat operator sequence and that thiosulfinate exposure causes de-repression dependent on a conserved cysteine residue, specifically Cys100. We further demonstrated that AltR controls expression of other *alt* genes and that mutations impairing AtlR de-repression reduce bacterial colonization on onion scales, highlighting the importance of AltR-mediated regulation in the adaptation of *P. ananatis* to onion-derived chemical defenses.

## Results

### AltR represses its own promoter and is de-repressed in response to allicin

AltR is predicted to be a TetR-family repressor and related to NemR ^30,31^. NemR repressed its own expression and the co-cistrionic *nemA* reductase gene by binding to an inverted repeat operator in the *nemR* promoter region ^40^. Inspection of the region upstream of *altR* in *P. ananatis* PNA 97-1R revealed a perfect inverted repeat motif (CAATCTAC [N6] GTAGATTG) partially overlapping the predicted -10 box and ribosome-binding site (RBS) of a putative σ70 promoter (Figure 1A). To build an *altR* promoter reporter, we inserted the full intergenic sequence between *altR* and *altG* (P_altR_) from PNA 97-1R to drive an autobioluminescence Lux gene cassete (figure 1A) which was introduced into various PNA 97-1R genetic backgrounds via site-specific Tn*7* transposition ^41^.

**Figure 1.**
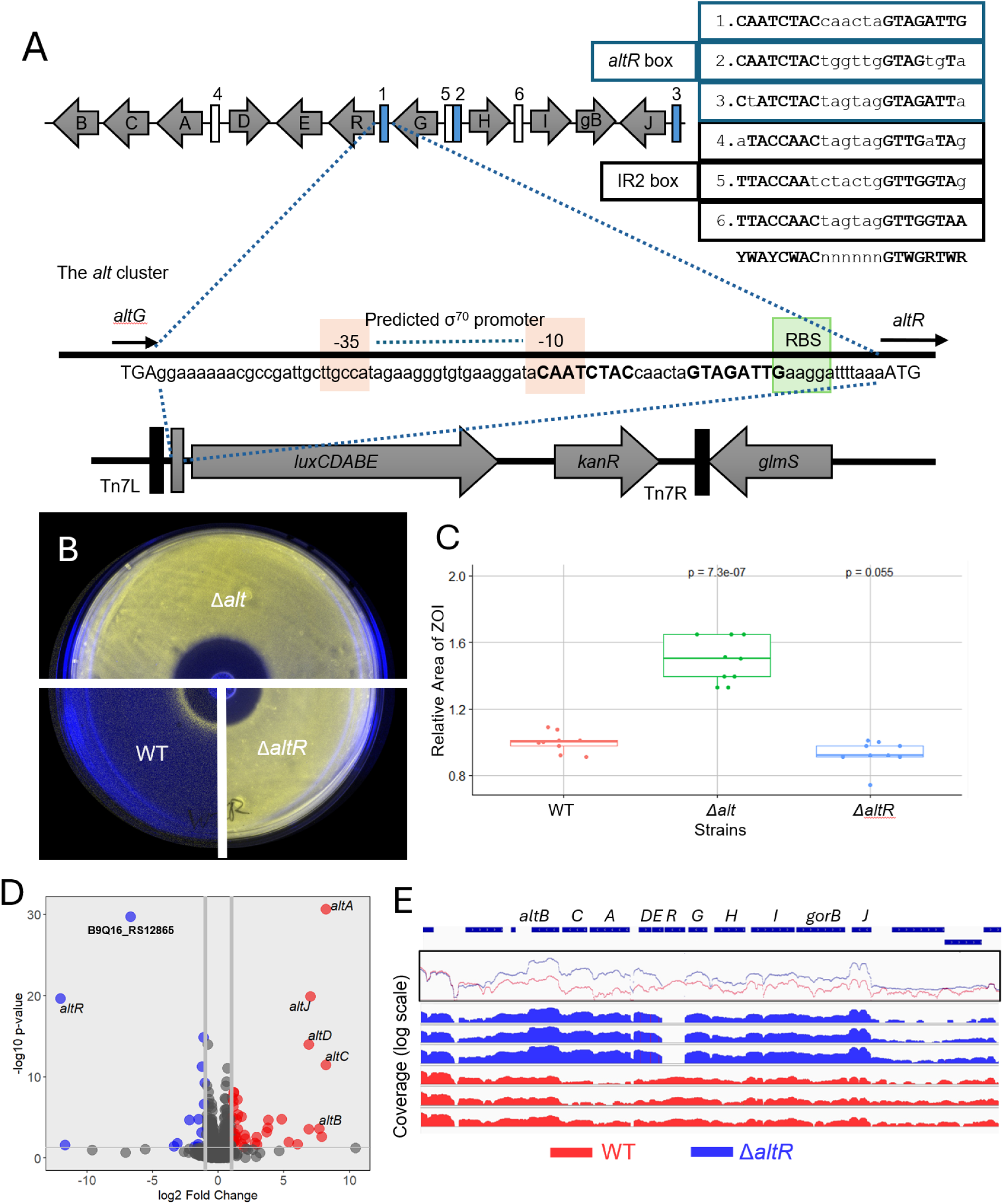
AltR represses its own promoter and the thiosulfinate-tolerance (*alt*) operon in *Pantoea ananatis*. **(A)** Schematic of the *altR* promoter bioluminescent reporter construct. The intergenic region between *altG* and *altR* was cloned as the *altR* promoter (*P_altR_*) upstream of the *luxCDABE* operon. Predicted AltR binding boxes and the inverted-repeat 2 [IR2] motif, the σ⁷⁰ promoter region (–35 / –10), and the ribosome-binding site (RBS) are indicated. Alignment of the predicted AltR and IR2 boxes is shown below. **(B)** Zone-of-inhibition (ZOI) assay showing *P_altR_LuxK6* activity in *P. ananatis* PNA 97-1R wild-type (WT), Δ*alt*, and Δ*altR* strains. The WT strain displays a distinct ring of bioluminescence at the ZOI border, indicating AltR de-repression in response to allicin exposure. **(C)** Quantification of relative ZOI area among WT, Δ*alt*, and Δ*altR* strains. Values represent mean ± SD of three biological replicates each with three technical replicates. Statistical significance was determined by one-way ANOVA followed by Tukey’s post hoc test (**p < 0.001**). The Δ*alt* mutant exhibited a significantly larger inhibition zone than WT, whereas Δ*altR* did not differ significantly from WT. **(D)** Volcano plot of differential gene expression from RNA-seq comparing WT and Δ*altR* strains (three biological replicates each). Genes that are up-regulated are highlighted in red, while down regulated genes are blue, with specific significant genes labelled. Deletion of *altR* resulted in strong up-regulation of *altA–altJ* and down-regulation of the YchH-like gene (B9Q16_RS12865). **(E)** Normalized RNA-seq read-coverage maps across the *alt* gene cluster in WT (red) and Δ*altR* (blue). Plots show mean log-scaled coverage for three biological replicates, with individual replicate tracks below. In Δ*altR*, coverage increases uniformly across *altA–altJ*, confirming AltR-mediated repression of the operon.

To test whether AltR regulates its own promoter, the bioluminescent reporter (P_altR_LuxK6) was introduced into wild type, Δ*alt*, and Δ*altR* backgrounds. Promoter activity was visualized in an allicin zone-of-inhibition (ZOI) assay (Figure 1B). Both Δ*alt* and Δ*altR* strains of PNA 97-1R displayed constitutive bioluminescence across the entire plate, consistent with loss of *altR*-mediated repression. In contrast, the wild-type strain exhibited a distinct ring of bioluminescence at the periphery of the inhibition zone, indicating promoter de-repression in response to allicin.

Quantification of inhibition zones (Figure 1C) showed that the Δ*alt* mutant had a significantly larger ZOI than either the wild type or Δ*altR*, confirming increased allicin sensitivity when the *alt* cluster is deleted. The Δ*alt*R strain displayed ZOI diameters similar to wild type, demonstrating that *alt*R deletion alone does not alter allicin sensitivity. The difference in Δ*alt* ZOI diameter also roughly correlated with region of bioluminescence de-repression in the WT strain indicating the AltR de-repression correlated with allicin tolerance. Overall, these observations indicate that AltR represses its own promoter under non-stress conditions and becomes de-repressed during allicin exposure.

### AltR represses the thiosulfinate tolerance (*alt*) cluster

To identify genes regulated by AltR, we performed RNA-seq analysis comparing the wild type and Δ*altR* strains of *P. ananatis* PNA 97-1R grown under rich media conditions (Figure 1D and 1E). Differential-expression analysis revealed that deletion of *altR* resulted in strong, coordinated upregulation of the *alt* cluster genes. The genes *altA*, *altB*, *altC*, *altD*, and *altJ* showed the highest induction, with log₂ fold-changes ranging from +5 to +10 (≈ 30–1000-fold increases).

Interestingly, a single gene outside the *alt* cluster, B9Q16_RS12865, encoding a predicted YchH-like protein, was the most strongly downregulated transcript in the Δ*altR* mutant (Figure 1D). Homologs of YchH in *E. coli* have been associated with stress-response modulation and redox balance ^42–44^, potentially suggesting that AltR may also influence broader oxidative-stress network in *P. ananatis*. No other loci showed substnatial transcriptional changes, indicating that AltR primarily functions as an *alt* cluster-specific repressor rather than as a global regulator.

Visualization of normalized read coverage across the *alt* locus (Figure 1E) further confirmed these transcriptional patterns. In wild-type cells, RNA-seq reads across *altA–altJ* were limited, consistent with promoter repression, whereas in Δ*altR* they increased uniformly across the cluster, reflecting de-repression of the cluster. Coverage dropped sharply within the *altR* coding region itself, validating the deletion.

Together, these data demonstrate that AltR primarily functions as a cluster-specific transcriptional repressor that governs expression of the thiosulfinate-tolerance (*alt*) operon. The strong downregulation of the YchH-like gene in the Δ*altR* mutant suggests that AltR activity may also indirectly influence additional stress-response pathways.

### Thiosulfinates specifically mediate de-repression of the *altR* promoter

Thiosulfinates are highly reactive organosulfur compounds that oxidize protein thiols mimicking certain aspects of oxidative stress^20^. To test whether AltR responds broadly to thiol-reactive or oxidizing compounds, or specifically to thiosulfinates, we performedzone-of inhibition (ZOI) assays using the P_altR_LuxK6 reporter strain exposed to various electrophilic chemicals (Figure 2A-G).

**Figure 2.**
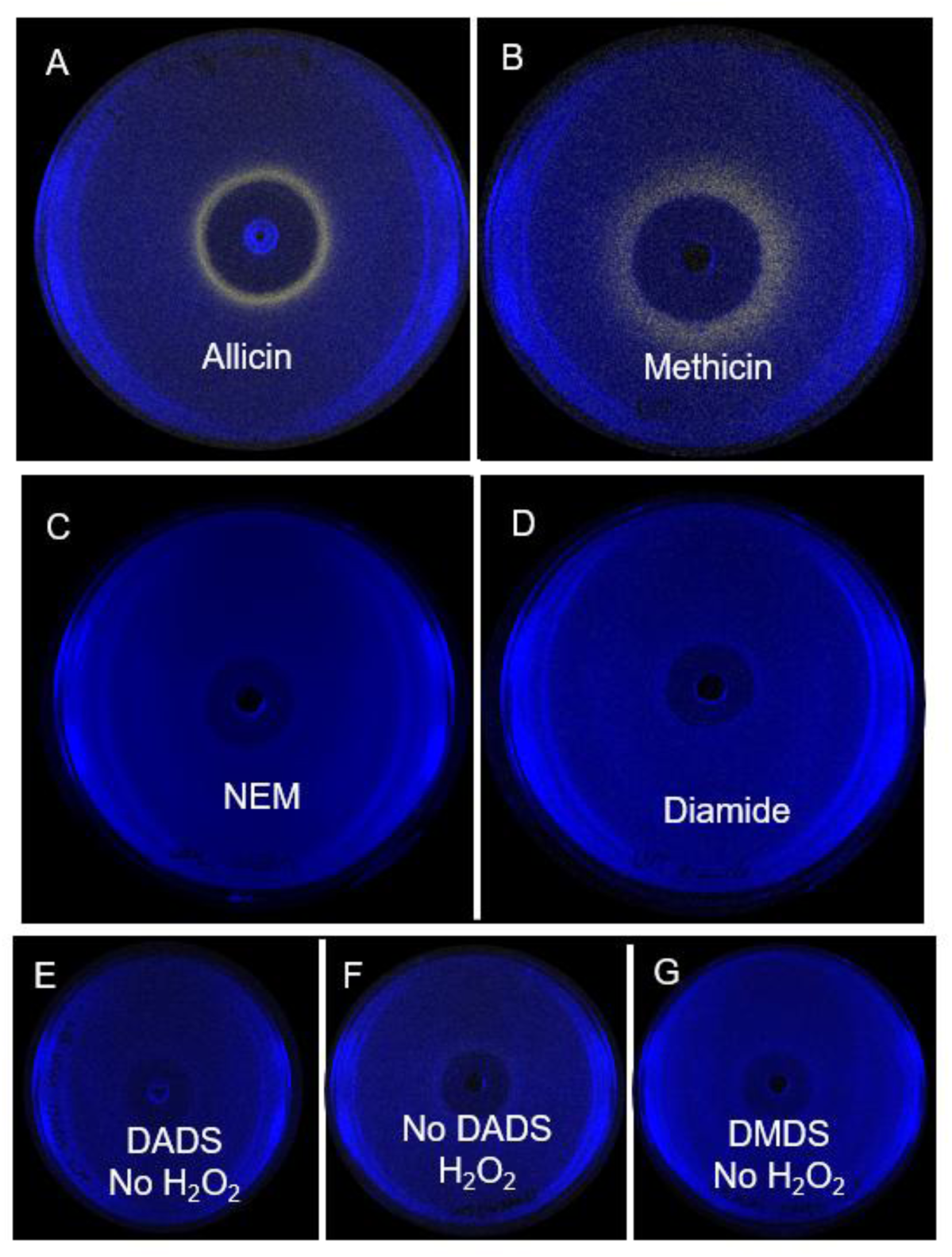
Thiosulfinates specifically trigger de-repression of the *altR* promoter in *Pantoea ananatis*. **(A–B)** Zone-of-inhibition (ZOI) assays showing *P_altR_::lux* reporter activity in *Pantoea ananatis* PNA 97-1R following treatment with the thiosulfinates allicin (A) and dimethyl thiosulfinate (“methicin”) (B). Both compounds induced a distinct ring of bioluminescence at the periphery of the inhibition zone, indicating AltR de-repression under sub-inhibitory concentrations. **(C–D)** Treatment with other thiol-reactive or oxidative compounds, including N-ethylmaleimide (NEM) (C) and diamide (D), no luminescent induction was observed, showing that AltR is unresponsive to these generic thiol oxidants. **(E–G)** The disulfide precursors diallyl disulfide (DADS) (E) and dimethyl disulfide (DMDS) (G), as well as hydrogen peroxide (H₂O₂) (F), also failed to trigger AltR de-repression. The absence of a luminescent ring in these treatments confirms that AltR activation is specific to thiosulfinates and not to disulfides or general oxidative stress.

As observed previously, allicin, the major thiosulfinate produced by garlic, induced a distinct ring of bioluminescence around the periphery of the inhibition zone (Figure 2A). A second thiosulfinate, dimethyl thiosulfinate (“methicin”), which is produced by both *Allium* and *Brassica* species ^45,46^, also triggered AltR de-repression, although the luminescence pattern appeared broader with less defined borders (Figure 2B).

In contrast, no promoter induction was observed in the presence of other thiol-reactive or oxidizing agents, indicating N-ethylmaleimide (NEM) and diamide, both known activators of the related NemR regulator in *E. coli* ^40,47^ (Figure 2C-D). Likewise, hydrogen peroxide (H₂O₂), a catalyst used in thiosulfinate synthesis, and the disulfide precursors diallyl disulfide (DADS) and dimethyl disulfide (DMDS) failed to elicit de-repression (Figure 2E–G).

Together, these results demonstrate that AltR specifically responds to thiosulfinates such as allicin and methicin, but not generally to oxidants or disulfides, confirming that AltR functions as a selective sensor of plant-derived thiosulfinates rather than a broad redox-stress regulator.

### AltR promoter de-repression coincides with tissue necrosis during onion infection

To investigate AltR-mediated regulation during onion tissue infection, we inoculated red onion scales with the *P. ananatis* PNA 97-1R P_altR_LuxK6 reporter strain to monitor both symptom development and bioluminescence over time (Figure 3). At 1 day post inoculation (dpi), a faint bioluminescence signal was detected at the inoculation site, corresponding to de-repression consistent with wound-based release of thiosulfinates. By 2 dpi, no visible bioluminescence was observed, suggesting limited atlR promoter activity during early asymptomatic colonization. At 3 dpi, the onset of necrotic lesion formation coincided with a pronounced increase in bioluminescence intensity at the infection site, indicating strong *altR* de-repression. By 4 dpi, as necrosis expanded, bioluminescence spread outward from the original inoculation point, mirroring the advancing zone of tissue collapse. These results demonstrate that *altR* promoter activation, and by extension, *alt* cluster expression, is tightly associated with host tissue necrosis, consistent with thiosulfinate production and bacterial adaptation to the environment of necrotic onion tissue.

**Figure 3.**
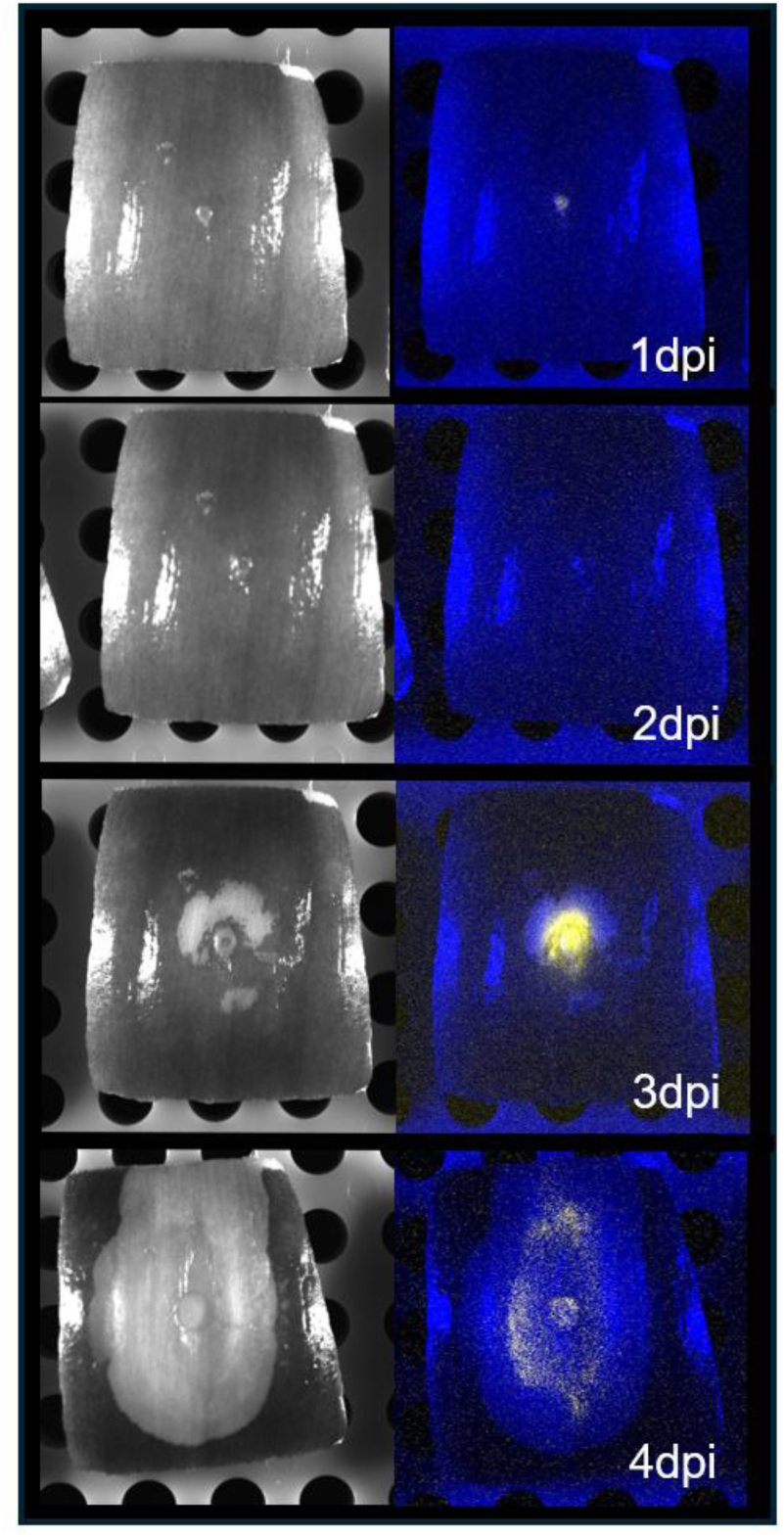
*altR* promoter de-repression coincides with necrosis during onion infection. Red onion scale assays were performed using bioluminescently labeled *Pantoea ananatis* PNA 97-1R *PaltRLuxK6* to visualize *altR* promoter activity during infection. Necrotic symptoms and corresponding bioluminescence (colored yellow) became evident at 3 days post-inoculation (dpi) and intensified at 4 dpi, coinciding with tissue collapse. The luminescence signal remained confined within necrotic tissue, consistent with thiosulfinate production from damaged onion cells. Note that signal intensities are not normalized across time points because images were captured independently on different days.

### The predicted AltR binding box is required for *altR* promotor repression

To determine whether AltR directly requires the predicted *altR* box inverted repeat (IR) sequence upstream of *altR*, we generated site-directed mutations within the putative operator region. Two mutant promoter constructs were designed: mod1, which introduced transversion mutations in one *altR* box half-site, and mod2, which altered both *altR* box half-sites. The resulting reporter strains (P_altRmod1_LuxK6 and P_altRmod2_LuxK6) were evaluated using ZOI assays. Both mutants lost the repression phenotype and exhibited display constitutive bioluminescence across the plate, indicating that integrity of the *altR* box sequence is required for AltR-mediated repression (Figure 4). This supports the prediction that the predicted *altR* box is required for AltR to regulate promoter activity. Interestingly, this specific IR motif (CAATCTAC[N6]GTAGATTG) is located only upstream of *altR, altG, altH* and *altJ*; but absent upstream of *altA* and *altD*, two of the most strongly upregulated genes in the Δ*altR* RNA-seq dataset (Figure 1A,1D and 1E). We identified an additonal inverted repeat (TTACCAAC[N6]GTTGGTAA), which we designated IR2, in the intergenic regions between *altA* - *altD*, *altG* - *altH*, and upstream *altI*. Given the strong upregulation of these genes in the absence of AltR, IR2 may represent an additional AltR operator site or a secondary cis-element involved in indirect regulation. Further biochemical studies are needed to determine whether AltR binds IR2 directly or through cooperative mechanisms involving other transcriptional regulators.

**Figure 4.**
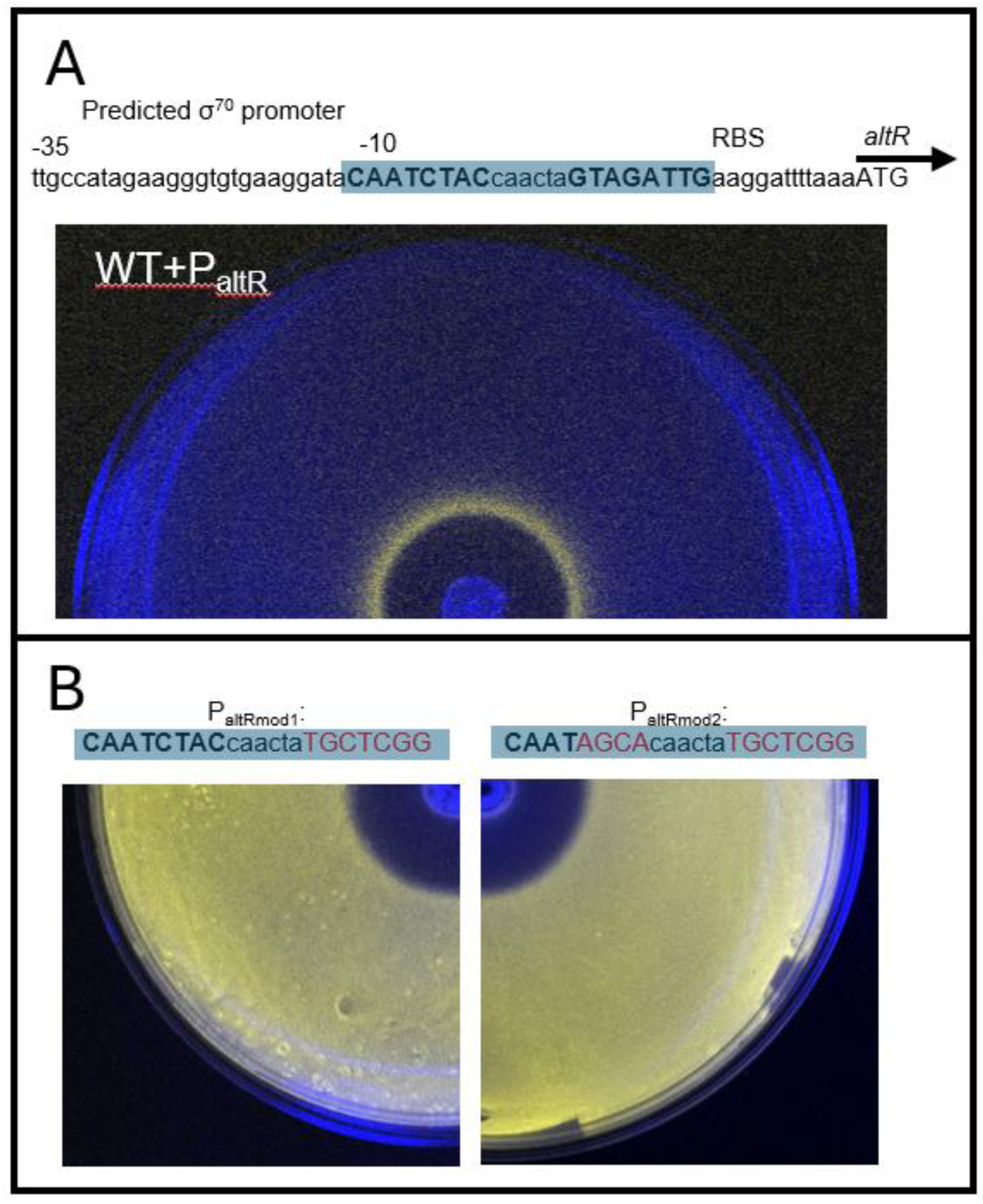
The predicted *altR* box is required for promoter repression. **(A)** Diagram of the *altR* promoter region showing the predicted *altR* box (CAATCTAC[N6]GTAGATTG). Inverted repeat (IR) half-sites are highlighted, with transversion mutations introduced in mod1 (one half-site) and mod2 (both half-sites). **(B)** Zone-of-inhibition (ZOI) assays of *P. ananatis* PNA 97-1R strains carrying P_altRmod1_LuxK6 and P_altRmod2_LuxK6 reporter constructs. Mutation of one or both IR half-sites led to constitutive bioluminescence across the plate, indicating complete loss of AltR-mediated repression.

### AltR cysteine residues are crucial for thiosulfinate-mediated de-repression

Because thiosulfinates are known to react directly with protein cysteine thiols ^26^ , we hypothesized that AltR cysteines would be crucial for sensing and responding to these compounds. The *P. ananatis* PNA 97-1R AltR protein consists of three cysteine residues: Cys100, Cys101, and Cys159. To test their roles in regulation, we constructed a complete panel of seven cysteine-to-serine substitution mutations, encompassing all combinations. In these mutants, serine replaces cysteine’s reactive thiol with a hydroxyl group. Each mutant alleles, designated according to residue substitution (e.g. AltR(C100S) = AltRSCC) was introduced into the *alt* cluster via allelic exchange. In allicin ZOI assays, the four alleles carrying the C100S substitution lost the allicin-induced de-repression phenotype (Figure 5A) and exhibited larger inhibition zones relative to wild type (Figure 5B), indicating increased sensitivity to allicin. Consistent results were observed with methicin treatments (Figure 5C-D). Mutations at Cys101 or Cys159 impacted basal or de-repressed bioluminescence levels, suggesting that these residues fine-tune AltR repression strength or inducibility rather than acting as the primary sensing site. Together, these findings indicate that Cys100 is essential for AltR-mediated thiosulfinate sensing, while Cys101 and Cys159 modulate regulation.

**Figure 5.**
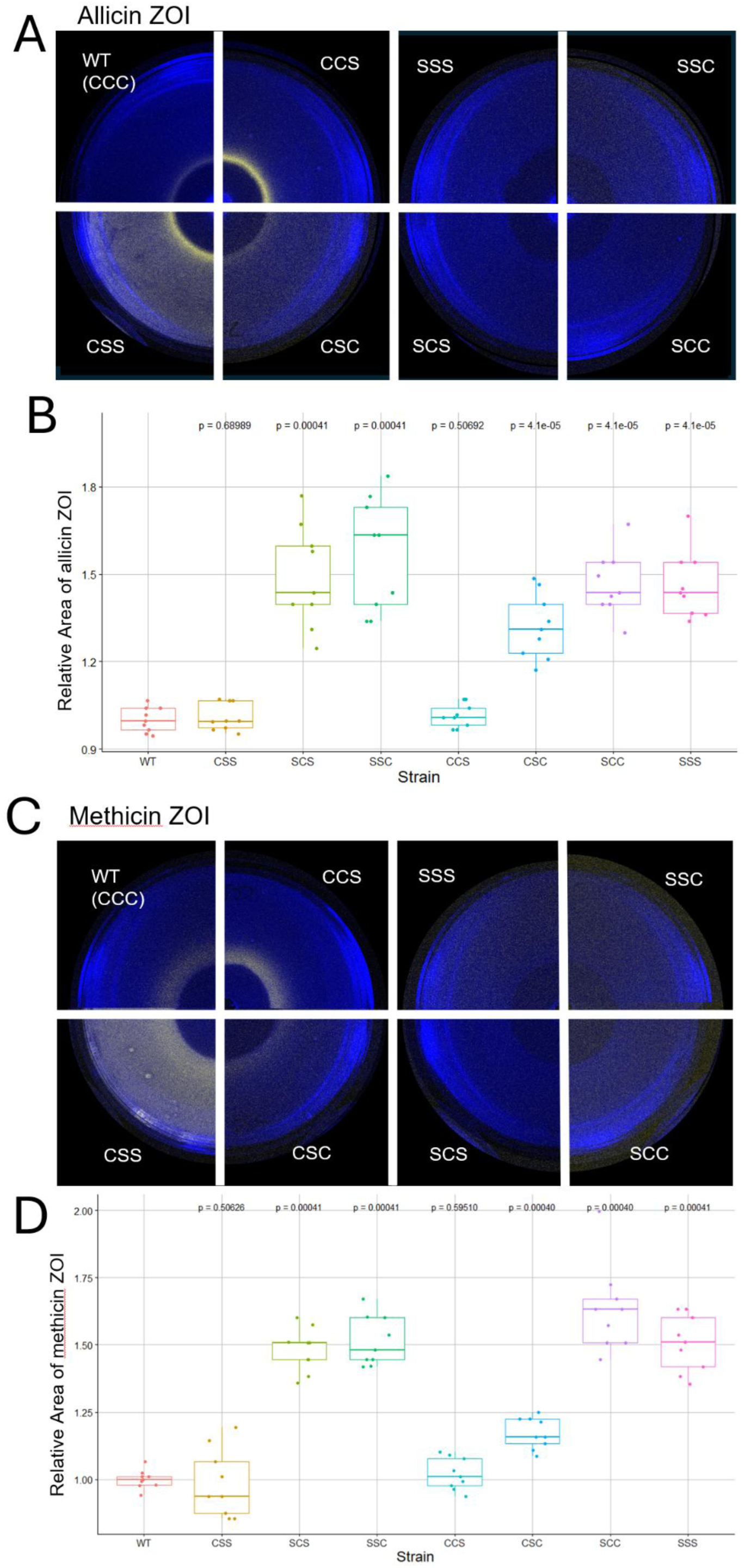
Cysteine residues of AltR are required for thiosulfinate-dependent de-repression. **(A)** Zone-of-inhibition (ZOI) assays showing bioluminescence from PNA 97-1R *PaltRLuxK6* strains carrying different *altR* cysteine-to-serine substitution alleles following treatment with synthesized allicin. Strains retaining Cys100 (e.g., CCC, CSS, CSC, SCC) displayed a luminescent induction ring at the edge of the inhibition zone, whereas all Ser100 (C100S) mutants lacked induction, indicating loss of AltR responsiveness to allicin. **(B)** Quantitative analysis of relative ZOI area for wild-type and mutant strains in response to allicin. Data represents three biological replicates, each with three technical replicates, analyzed by one-way ANOVA followed by Tukey’s multiple-comparison test (*p* < 0.001). **(C)** ZOI assays of the same *altR* cysteine mutants exposed to methicin (dimethyl thiosulfinate). As with allicin, only strains retaining Cys100 exhibited promoter induction, confirming that this residue mediates AltR redox sensing of thiosulfinates. **(D)** Relative ZOI area measurements for methicin assays analyzed as described in (B).

### Quantification of de-repression and leaky repression in response to thiosulfinates

To quantitatively assess AltR-mediated promoter activity, we used a Newton 7.0 bioluminescence detection system (Vilber Smart Imaging) capable of photon-count quantification. ZOI assay images were analyzed by six regions of interest (ROIs) within both the inhibition zone and the background region, and the maximum photon counts were recorded for each ZOI (Figure 6A).

**Figure 6.**
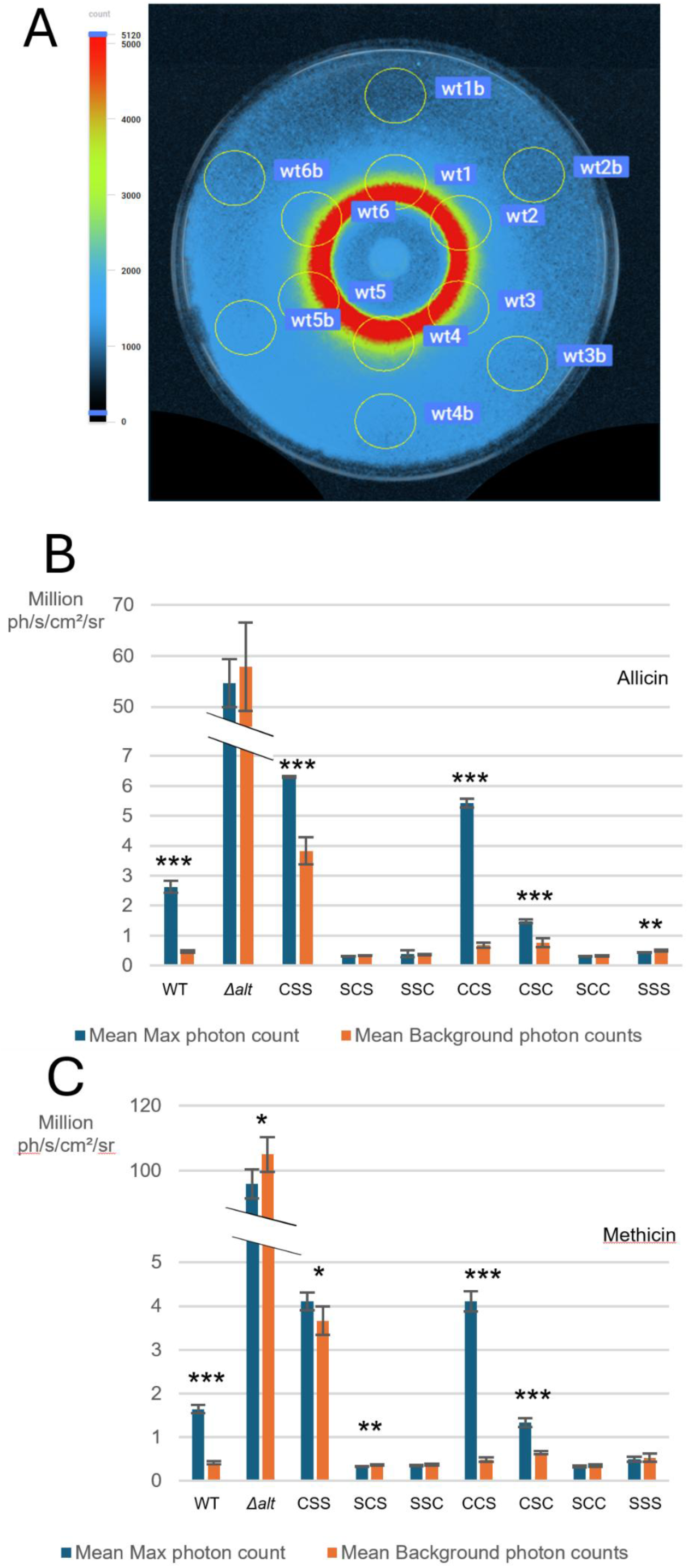
Quantitative analysis of cysteine-to-serine substitutions on *AltR*-mediated de-repression and leaky repression in response to thiosulfinates. **(A)** Zone-of-inhibition (ZOI) assay imaged using the Newton 7.0 (Vilber Smart Imaging) bioluminescence detection system. Six regions of interest (ROIs) were sampled from both the induction ring and the background region to measure photon counts, as illustrated. **(B)** Quantification of bioluminescence for *P. ananatis P_altR_LuxK6 altR* cysteine mutants exposed to allicin. Mean maximum photon counts (blue bars) within the inhibition zone and mean background photon counts (orange bars) are shown for each allele. Strains retaining Cys100 exhibited significant de-repression compared to their background levels, while C100S mutant alleles did not. **(C)** Quantification of bioluminescence for the same mutant panel exposed to methicin. Similar trends were observed, confirming that Cys100 is essential for thiosulfinate-dependent induction. Error bars represent standard deviation of three biological replicates, each with three technical replicates. Significance was determined by two-sample *t*-tests: *p* < 0.05 (\**), p < 0.01 (**), p < 0.001 (\*\*\**).

Bioluminescence from assays treated with allicin and methicin were quantified and plotted separately (Figure 6B-C). All strains retaining Cys100 exhibited significantly higher mean maximum photon counts within the inhibition zone compared to background regions, confirming AltR de-repression upon thiosulfinate exposure. In contrast, mean maximum photon counts of C100S mutant alleles are not higher than their background, consistent with their loss of responsiveness to thiosulfinates.

Notably, several AltR alleles exhibited distinct regulatory behaviors. The CSS allele showed elevated background bioluminescence, indicating leaky repression, while the CCS allele displayed stronger de-repression than wild type, suggesting enhanced promoter induction. Conversely, the CSC allele exhibited reduced de-repression compared to wild type, correlating with its weaker resistance phenotype observed in ZOI assays. These quantitative measurements reinforce that Cys100 is essential for thiosulfinate sensing, while Cys101 and Cys159 modulate AltR sensitivity and basal repression strength.

### AltR Cys mutations alter bacterial proliferation in onion tissue

To determine how *altR* cysteine mutants affect bacterial fitness during host colonization, we performed bacterial load assays on red onion scales using the complete set of *altR* cysteine-to-serine substitution strains. Because each mutant retains the HiVir pantaphos biosynthetic gene cluster, all were capable of producing necrotic lesions. After four days of infection, the CCS and CSS mutants reached bacterial population levels comparable to wild type, while the CSC mutant exhibited significantly lower population, similar to the Δ*alt* strain (Figure 7A). This result parallels the *in vitro* inhibition zone assays (Figure 5), where the CSC mutant displayed increased sensitivity to thiosulfinate compared with WT, CCS, and CSS. All four C100S-containing mutants exhibited markedly reduced bacterial loads relative to WT yet still achieved higher population than the Δ*alt* strain (Figure 7B). These findings demonstrate that Cys100 is critical for AltR function and thiosulfinate tolerance in vivo, and that specific cysteine combinations modulate pathogen proliferation within onion tissue.

**Figure 7.**
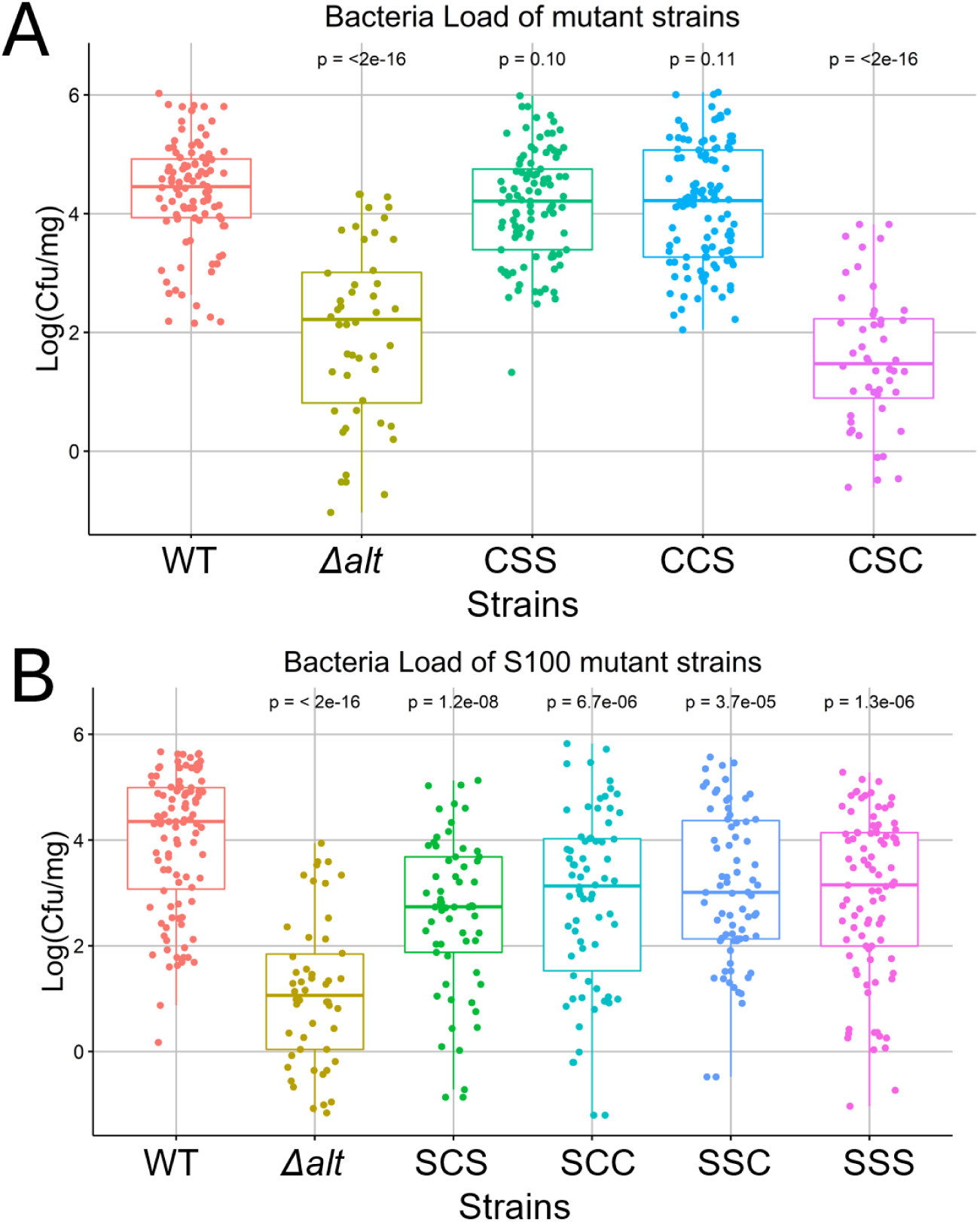
AltR cysteine mutations alter bacterial proliferation in onion tissue. **(A)** Bacterial population sizes of PNA 97-1R *PaltRLuxK6* strains carrying non-C100S *altR* alleles (CCC, CSS, CSC, CCS) on red onion scales at 4 days post-inoculation (dpi). The CSC mutant exhibited significantly reduced bacterial loads, comparable to the Δ*alt* strain. **(B)** Bacterial populations of *C100S* mutant strains (SCC, SCS, SSC, SSS) at 4 dpi. All C100S alleles showed decreased proliferation relative to wild type but maintained higher populations than Δ*alt*. Each data point represents a single biological replicate (six biological replicates, five technical repeats per strain). Data were log₁₀-transformed prior to analysis. Statistical significance was assessed using one-way ANOVA followed by Tukey’s multiple-comparison test (*p* < 0.001). Pairwise comparisons against the WT control were evaluated by *t*-test.

## Discussion

In this study we present evidence that the AltR functions in an analogous manner to the TetR-family repressor NemR in that it (1) represses its own transcription and (2) the transcription of corresponding stress tolerance factors and (3) is derepressed by the offending chemical stressor. In addition, AltR regulation requires an inverted repeat operator and is dependent on a key conserved cysteine residue ^40^. The deletion of *altR* results in increased transcription of the ten other genes in the PNA 97-1 *alt* gene cluster. The AltR-mediated repression of P_altR_ is relaxed by exposure to either allicin or methicin as well as *in vivo* in necrotic onion tissues where thiosulfinates would be natively produced. AltR de-repression is driven by thiosulfinates but not by other oxidizers nor disulfide derivatives. Lastly, mutational evidence supports for Cys100 being the key cysteine residue required for thiosulfinate-mediated de-repression, while Cys101 and C159 contribute to fine-tuning AltR-mediated regulation.

Previous work supported AltR functioning as a repressor of thiosulfinate tolerance ^31^. AltR repressed expression from the P_altR_ Lux reporter, but de-repression was observed at the border of an allicin thiosulfinate inhibition zones (figure 1B). Notably, the Δ*alt* strain had increased thiosulfinate sensitivity in the ZOI assay with the increased diameter correlating with the P_altR_ Lux “induction ring” of de-repression in the WT. This suggested that AltR-mediated de-repression coincides with increased phenotypic thiosulfinate tolerance. We also observed that *altR* promoter de-repression occurs *in vivo*, correlating with the onset of necrotic symptom where dead onion cells release thiosulfinates (figure 3). We identified a perfect inverted repeat (CAATCTAC [N6] GTAGATTC) in the *altR* promoter region as a potential *altR* box operator as well as two additional imperfect altR box operators in the *alt* cluster (figure 1A). We also identified a second inverted repeat (TTACCAAC [N6] GTTGGTAA) present in *alt* gene intergenic regions which we named Inverted Repeat 2 (IR2) also with two others imperfect IR2 boxes found in the *alt* cluster (Figure 1A). We provided genetic evidence that the predicted *altR* box within the *altR* promotor region is required for AltR-mediated repression of the P_altR_ Lux reporter. Mutating the sequence of the predicted AltR binding box results in loss of repression (figure 4).

In the absence of *altR*, all ten remaining *alt* genes were upregulated consistent with regulating thiosulfinate tolerance. The *altA, altB, altC, altD*, and *altJ* genes are some of the most up-regulated genes (figure 1D). However, only the *altJ* gene has a predicted upstream *altR* box. The expression of *altA, altB, altC,* and *altD*, may be mediated via the intergenic IR2 box either through direct AltR recognition of IR2 as a second operator or indirectly through AltR-regulation of an additional transcription factor. No other annotated transcription factors were upregulated in the *altR* mutant, however a truncated ORF (B9Q16_23175) predicted to encode a HtH motif upstream of *altB* was upregulated. The *altR* box and IR2 consensus do form an inverted repeat motif with several shared bases YWAYCWAC [N6] GTWGRTWR, thus AltR may recognize both operator sequences (Figure 1A). Determining AltR DNA binding specificity will be an important area for future study and may provide more insights into the unexpected behavior of the AltR CSC allele. Among the RNA seq results, a single gene was strongly down regulated, locus number B9Q16_RS12865. This gene is predicted to be a homolog of an *E coli ychH*, which is a gene of unknown function identified from one study to be regulated in response of hydrogen peroxide, cadmium, and acid stress ^44^.

Previous studies demonstrated that the TTG clusters of *Pseudomonas fluorescens* strain PfAR-1 confer tolerance to synthesized allicin but not to other oxidant compounds like H_2_O_2_, or NEM^32^. NemR responds to reactive electrophile compounds, e.g. NEM, bleach, and diamide, to regulate bacterial tolerance to this class of compounds. Thus, we reasoned that AltR would specifically respond to thiosulfinates in a similar manner. We tested the responsiveness of AltR against several oxidative compounds. NEM and diamide were chosen to test because NemR responds to both NEM and diamide, and diamide, like thiosulfinates, oxidizes thiols (figure2C, 2D) ^32,40^. AltR de-repression occurred in response to allicin and methicin but not to NEM, diamide, H_2_O_2_ or the allyl-disulfide or methyl-disulfide precursors used for thiosulfinate synthesis (figure 2E, 2F, 2G). Thus, AltR has some degree of specificity for thiosulfinates.

Thiosulfinates vary based on their side chains ^24,48,49^ although they share the same functional group and can participate in similar chemical reactions. Allicin is the characteristic thiosulfinate of garlic and functionally there is little to no production of allicin by damaged onion cells. Onions produced a mixure of different kinds of symmetric and asymmetric thiosulfinates derived from either isoalliin, methiin, or propiin^20^. This is why we included synthesized methicin which can be produced by both onions and garlic as well as by *Brassica spp*. We observed similar overall responses to both allicin and methicin exposure.

PNA 97-1 AltR encodes three cysteines, Cys100 Cys101, and Cys159. However, in a sequence alignment of the *Pseudomonas* and *Pantoea* AltR, only Cys100 was conserved. We generated Cys to Ser mutants for all possible combinations of the Cysteine residues to see how the activity of the AltR repressor was impacted. After creating the 7 alleles of Cys to Ser and performing inhibition zone assays, we learned that Cys100 is, indeed, the most critical for thiosulfinate-mediated de-repression as only strains with this residue retain the thiosulfinate induction ring phenotype. Strains without this C100S mutation also all show increased sensitivity towards thiosulfinates comparable to a WT strain, implying AltR with the C100S stays bound to the altR box even in the presence of thiosulfinates. This behavior was consistent result for both allicin and methicin (figure 5).

Quantitative photon counts for the bioluminescence signal in the induction ring and background showed similar results although also demonstrated that Cys101 and Cys 159 tune both in the intensity of de-repression and the strength of repression (figure 6). For instance, the C159S allele has exaggerated de-repression resulting in a brighter induction ring than that produced by the WT AltR allele, whereas the C101S/C159S double mutant has an elevated background Lux expression consistent with leaky AltR repression. It is also interesting to note that the CSC mutant has higher thiosulfinate sensitivity compared to other C100S strains (Fig 5B and 5D) and reduced bacterial load in onion scales after 4 days post inoculation (figure 7). We are, as of yet, unclear how Cys101 mediates a more dramatic impact on bacterial fitness in onion tissue than the four thiosulfinate non-responsive C100S alleles.

In conclusion, this study provides new mechanistic insights into the regulation of the allicin tolerance gene cluster. As failure of AltR to derepress is associated with reduced bacterial loads in onion tissue, identification of antagonist compounds that can interfere with the capacity of AltR to respond to thiosulfinates may have utility both for management of onion center rot disease and potentially other diseases of onion caused by bacteria.

## Acknowledgements

This project was supported by USDA-NIFA-OREI 2023-51300-40913 to B. Dutta and B. H. Kvitko, and HATCH project 7002999 from the USDA National Institute of Food and Agriculture to B. H. Kvitko. Dr. Jovana Mijatovic helped part of the data analysis by sharing her R codes.

## Author contributions

B. H. Kvitko conceptualized this research. B. H. Kvitko and B. Dutta supervised this research. H. Jan performed the experiments and analyzed the data. F. Kong and L. Yang contributed to the RNA seq analysis. H. Jan, F. Kong, M. P. MacLellan and B. H. Kvitko wrote and revised the manuscript. All authors read, edited and approved of the final manuscript.

## Material and Methods

### Bacterial strains and cultural conditions

Overnight cultures of *Escherichia coli* and *Pantoea ananatis* were routinely cultured from single colonies recovered on LB parent plates and were grown in 5 mL of LB medium in 14 mL culture tubes placed in incubator of 28°C (*P. ananatis*) or 37°C (*E. coli*) with 200rpm shaking 18-22 h. See Table S1 for list of strains and plasmids.

### Bioluminescence signal imaging

Imaging was performed with the lab imager (analyticJena UVChemStudio). For strong bioluminescence signal strains (Mostly Δ*alt* strains), 2-minute exposure time was used; for weak bioluminescence strains, 5-minute exposure time was used. Strong signal strains and weak signal strains were not imaged together because the weak signal strains were overshadowed. An image under white light was also taken for each bioluminescence image for generating image overlays with ImageJ, with the non-bioluminescence background colored blue and the bioluminescence signal colored yellow.

### Construction of the pTn7P_altR_LuxK6 vector

The bacterial promoter P_altR_, intergenic region of *P. ananatis altG* and *altR*, was synthesized as a dsDNA gblock by IDT (Table S4.) and cloned into the StuI and DraIII double*-*digested backbone of pTn5/7LuxK6 ^41^ via Gibson assembly (New England Biolab). The insertion of the P_altR_ promoter was confirmed by sequencing.

### Mini-Tn7 labelling of strains

*E. coli* RHO3 pTNS3, *E. coli* RHO5 pTn7P_altR_LuxK6, and the target *Pantoea* strain were combined in a tri-parental mating ^50,51^. 5 mL LB cultures of each strain were grown overnight. The following day 1 mL of each culture was pelleted and resuspended in 100 mL of fresh LB. 10 μL of each concentrated culture was added to a clean tube. LB DAP plates were prepared and sterile nitrocellulose membranes were placed on the plates. 30 μL droplets of the mixed cultures, and independent strain controls were placed on the nitrocellulose membranes to dry. Following overnight incubation, the mixture was removed from the nitrocellulose membrane using a sterile loop and resuspended in 1 mL LB. 200 μL of the incubated mixed culture suspension was plated on Km LB (with no DAP) selection plates. The following day Km resistant colonies were selected confirmed for luminescence with 2 m exposure settings on the lab imager (analyticJena UVChemStudio).

### Preparation of allicin and methicin stock solutions

Fresh allicin and methicin stock solutions were prepared for experiments using a modified microscale synthesis protocol ^52^. Solutions were stored at -20 C and used within three days of prep. For allicin synthesis, mix 5 μL of diallyl disulfide (DADS) 96% (Carbosynth), 25 μL of glacial acetic acid, and 15μL of 30% hydrogen peroxide were added to 4x 200μL PCR tubes. For methicin synthesis, dimethyl disulfide 99% (Sigma) was used in the same way. Tubes were agitated in a 28°C shaker for at least 4h. Following agitation, the reaction in all 4 tubes were quenched in 1 mL of methanol. This methanol allicin mixture was used as the stock synthesized allicin preparation and was used directly in ZOI assays. Controls for disulfide precursors testing (figure 2) were made following the same protocol with the absence of hydrogen peroxide.

### Zone of Inhibition assay

Styrene Petri plates (100 x 15 mm) with 20 mL LB adding the required antibiotics were spread with 400μL of overnight bacterial culture using a sterile cotton swab. A 0.125 cm^2^ agar plug was removed from the center of the plate with a biopsy punch. 50 μL of treatment solution was added into the agar well. Treatment solutions that were used in this research include the allicin or methicin stocks, 30 mM NEM 99% (Thermoscientific) dissolved in ethanol, and 150 mM diamide 97% (Tokyo Chemical Industry) dissolved in DMSO. Plates were incubated for 24 h at RT and evaluated for a zone of inhibition (cm^2^) by measuring the diameter, calculating the relative inhibition area for statistical analysis. Raw data was processed through Excel (Microsoft), and Rstudio (Package agricolae) was used for conducting one-way ANOVA analysis and t-tests, as well as drawing graphs.

Images of the plates were taken via the lab imager (analyticJena UVChemStudio) with 2 or 5 minutes exposure (depending on the strength of bioluminescence) for bioluminescence signals and a normal condition image under white light for overlapping, color coded bioluminescence as yellow and white light image as blue in the images.

### Red onion scale necrosis assay

Consumer produce red onions (*Allium cepa*. L., red onion) were purchased, cut to approximately 3 cm wide scales, sterilized in a 3% household bleach solution for 20 seconds, then rinsed in dH_2_O for 4∼5 times gently. Red onion scales at least 0.5cm thick with a healthy undamaged appearance were used. Scales were placed in a potting tray (27 x 52 cm) containing two layers of paper towels pre-moistened with 90 mL of distilled water. The plastic removable portion of 20 μL pipette trays (sanitized before use) were positioned on top to prevent direct contact between the paper towels and onion scales. Individual onion scales were wounded cleanly through the scale with a sterile 20 μL pipette tip and inoculated with a 10 μL drop of bacterial culture of 0.3 OD600. Sterile deionized water was used as a negative control inoculum. The tray was covered with a plastic humidity dome and incubated at RT for 96 h; images were taken every 24 h with the lab imager (analyticJena UVChemStudio) to record the disease development.

Quantification of bacteria growth (cfu/mg) within onion scales was performed after red onion scales assays by sampling small chunks of tissues (approximately 50∼100 mg) 1cm away from the inoculation site. The sampled tissues were weighed and placed in plastic maceration tubes with 200 mL of sterile dH_2_O and three Xmm diameter high density zircon beads. The tubes beat at 4 m/s for 1 min using the Bead Ruptor Elite Bead Mill Homonizer (Omni International). Samples then underwent 10-fold serial dilution (20ul to 180 ul) with sterile water up to 10^-7 in a sterile 96 well plate. 10ul of the diluted suspensions were then plated on LB Rif plates, incubated overnight in 28°C. Colonies were counted the next day, normalized by sample weight to quantify bacteria load. Raw data was processed through Excel (Microsoft), and Rstudio (Package agricolae) was used for conducting one-way ANOVA analysis and t-tests, as well as drawing graphs.

### RNA seq analysis

Total RNA was extracted by Quiagen miRNeasy Mini kit from overnight culture of PNA 97-1R PaltRLuxK6 as well as a Δ*altR* version of the same strain. The RNA extract of 3 biological repeats of each strain were sent to Azenta Life Science to undergo DNAse treatment and next generation sequencing for RNA seq purpose. The raw reads were checked for quality control using FastQC and trimmed by Trimmomatic. The clean RNA-seq reads were aligned against the *Pantoea ananatis PNA 97-1R* reference genome. Stringtie was used to assemble into transcripts according to the *Pantoea ananatis PNA 97-1R* annotation. DEseq2 was used to analyze differential expression analysis. Volcano plot was made in R. Deeptools and bedtools were used to calculate the mean coverage of three biological replicates and Integrative Genomics browser was used to virtualize the coverage map.

### AltR box modification construction

Mutated variants of the *altG* and *altR* intergenic region were synthesized as dsDNA gblocks by IDT (Table S3) and cloned into the StuI and DraIII double*-*digested backbone of pTn5/7LuxK6 ^41^ via Gibson assembly. The insertion of the two mod box promoter was confirmed by sequencing. After assembly of the plasmid, the modified *altR* box were inserted into PNA 97-1R with the mini-Tn7Lux labelling method mentioned in previous sections, making the two strains PNA 97-1R P_altRmod1_LuxK6 and PNA 97-1R P_altRmod2_LuxK6.

### Construction of cysteine mutants of *altR* expression vectors and allelic exchange of cysteine to serine mutants

Cysteine residue mutants of *altR* were built in gateway expression vectors and later transformed in PNA 97-1R Δ*altR* PaltRLuxK6 for plasmid-based complementation. The pDONR221::altR plasmid was made by PCR amplifying the altR gene region then insertion into the pDONR221 empty plasmid with BP cloning. See Table S2 for primer sequences. PCR was conducted under the following condition: 95℃ for 30s, then repeat 35 cycles of 95℃ denaturing for 30s, 60℃ annealing for 30s, 72℃ extension for 1m. Final extension at 72℃ for 4m. The *altR* mutants, SSS, CSS, SCS were synthesized by Genscript in the Gateway compatible plasmid pUC57 with an additional BspE1 site. The four other combinations of the cysteine mutants, CSC, CCS, SSC, SCC, were made by restriction enzyme BspE1 to swap the C-terminal portions including *altR* Cys_159_ from the *altR* sequence, then ligation with T4 DNA ligase (ThermoScientific) was performed.

The flanks around altR(100-159) were first synthesized along with a dual BbsI cutsite in the middle, sequence named attB-altRd100∼159-flanks-BbsI (Table S4), then was inserted into allelic exchange vector pR6KT2G via BP clonase recombination. The mutated AltR(100∼159) regions were then amplified with PCR by using the expression vector mutants as templates under the following PCR condition: 95℃ for 30s, then repeat 35 cycles of 95℃ denaturing for 30s, 60℃ annealing for 30s, 72℃ extension for 1m. Final extension at 72℃ for 4m. Gibson assembly was then utilized to assemble 7 different allelic exchange plasmids that includes all 7 different combinations of cysteine to serine mutants (pR6KT2G_altRXXX). The 7 assembled plasmids were transformed into donor strain RHO5, then introduced into target strain PNA 97-1R Δ*altR*(100-159) P_altR_LuxK6 by bacterial mating on LB DAP plates. Merodiploids were recovered 24 hours later on LB Gm plates. Two merodiploid colonies of the same mutant were then selected and added in a tube of 1mL sterile LB with 3mL sterile 1M Sucrose to grow for 24hr at 37°C. Counter selection was done the next day by plating 1×10^5^ or 1×10^6^ dilution of the culture on LB Xgluc plates. Colonies that are shown blue has not lost the inserted gateway plasmid, therefore white colonies were patch plated to find loss of Gm resistant. Altered bioluminescence were also checked for the newly made strains. Further sequencing were performed to confirm the allelic exchanged mutants.

### Photon quantification

Bioluminescence photon count data images were taken by Newton 7.0 (Vilber Smart Imaging) under software auto recommendation settings. Images were then processed through Kaunt 2.5 (Vilber Smart Imaging) and areas of interest were selected through it (figure 6A). The Max ph/s/cm²/sr data for regions of interests were then exported to Excel (Microsoft) and processed to draw graphs. Rstudio (Package agricolae) was used for conducting t-tests.

---

**Table S1.**
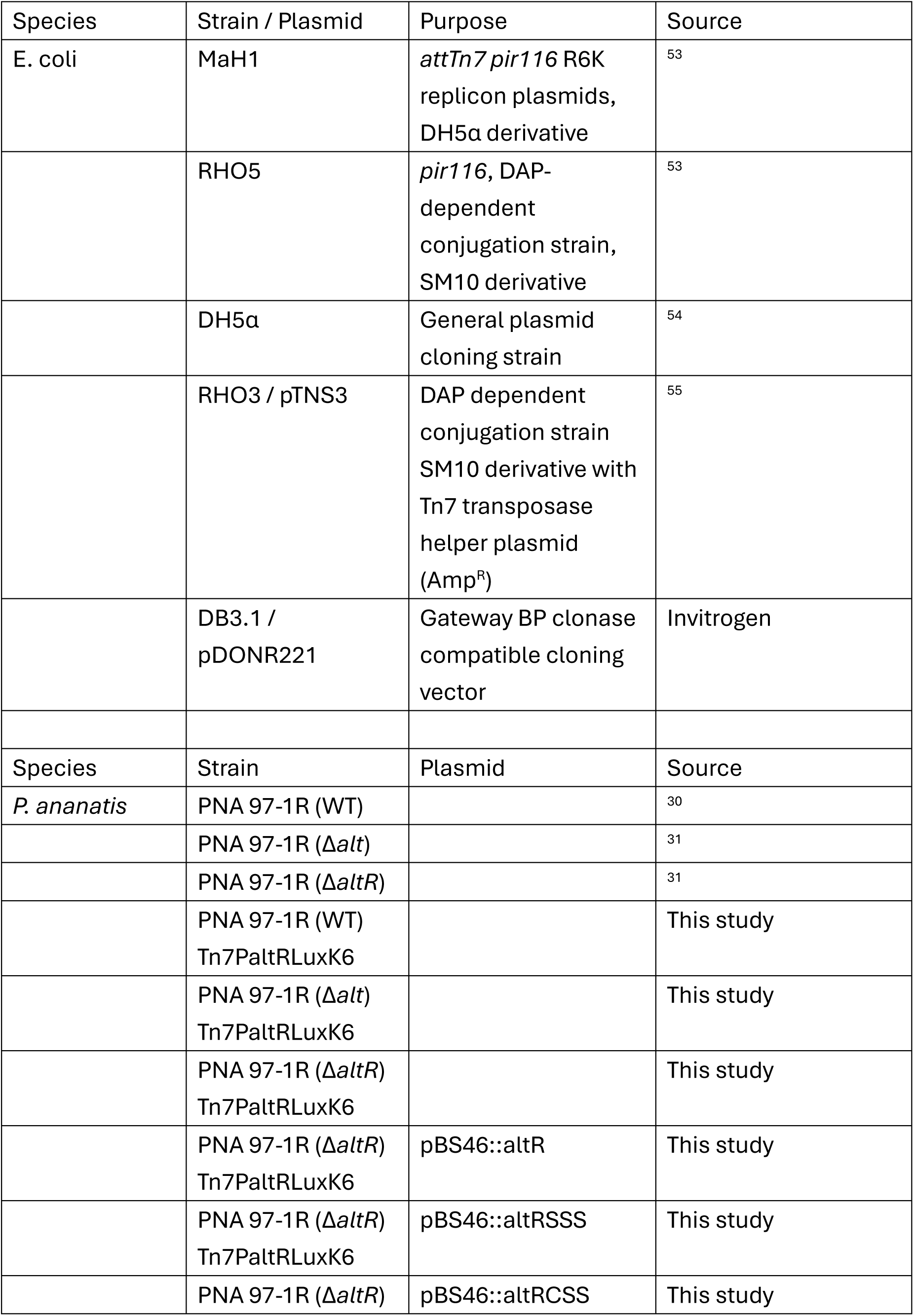
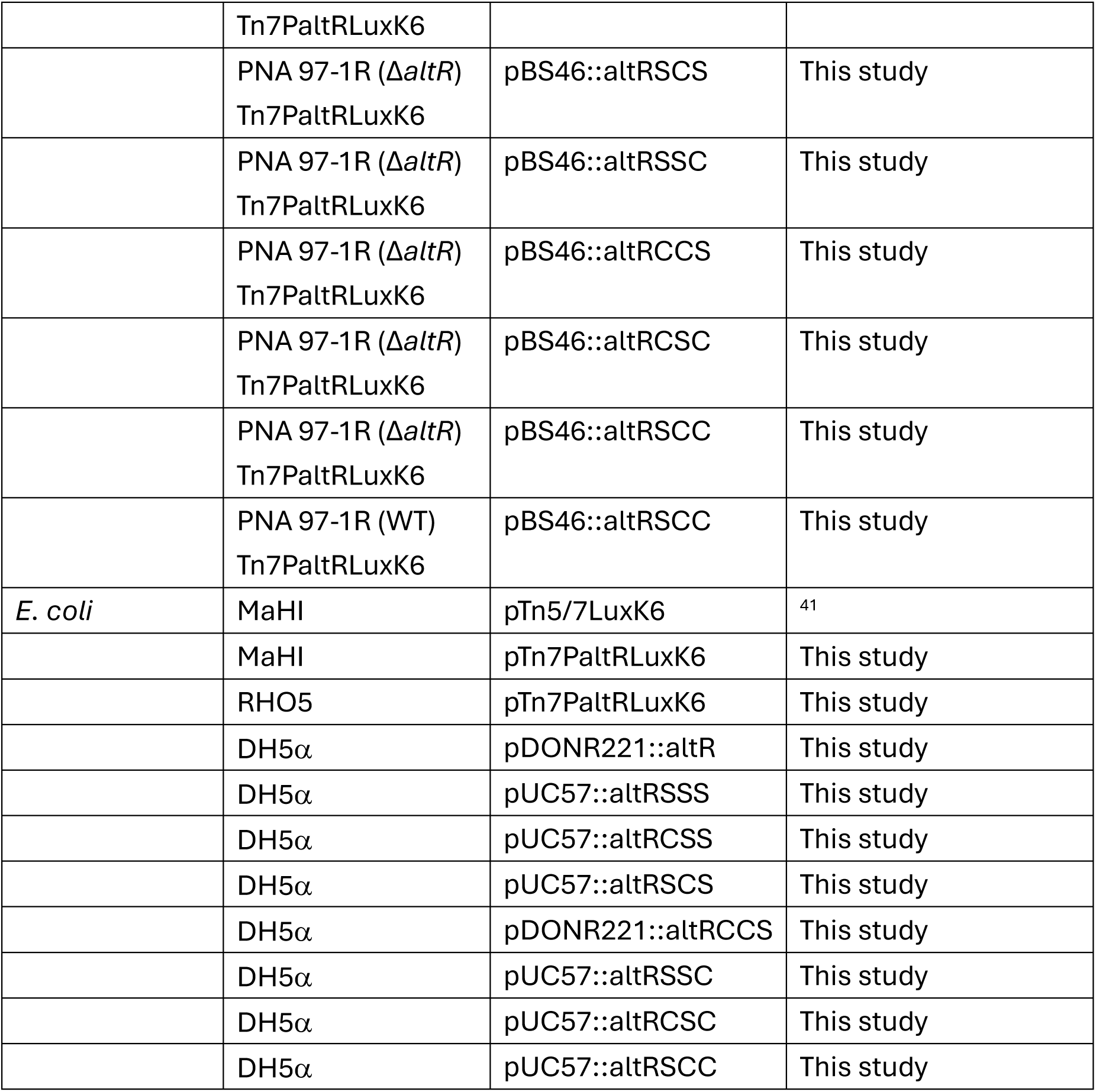
General cloning, deletion, and labeling strains and vectors used in this study.

**Table S2.**
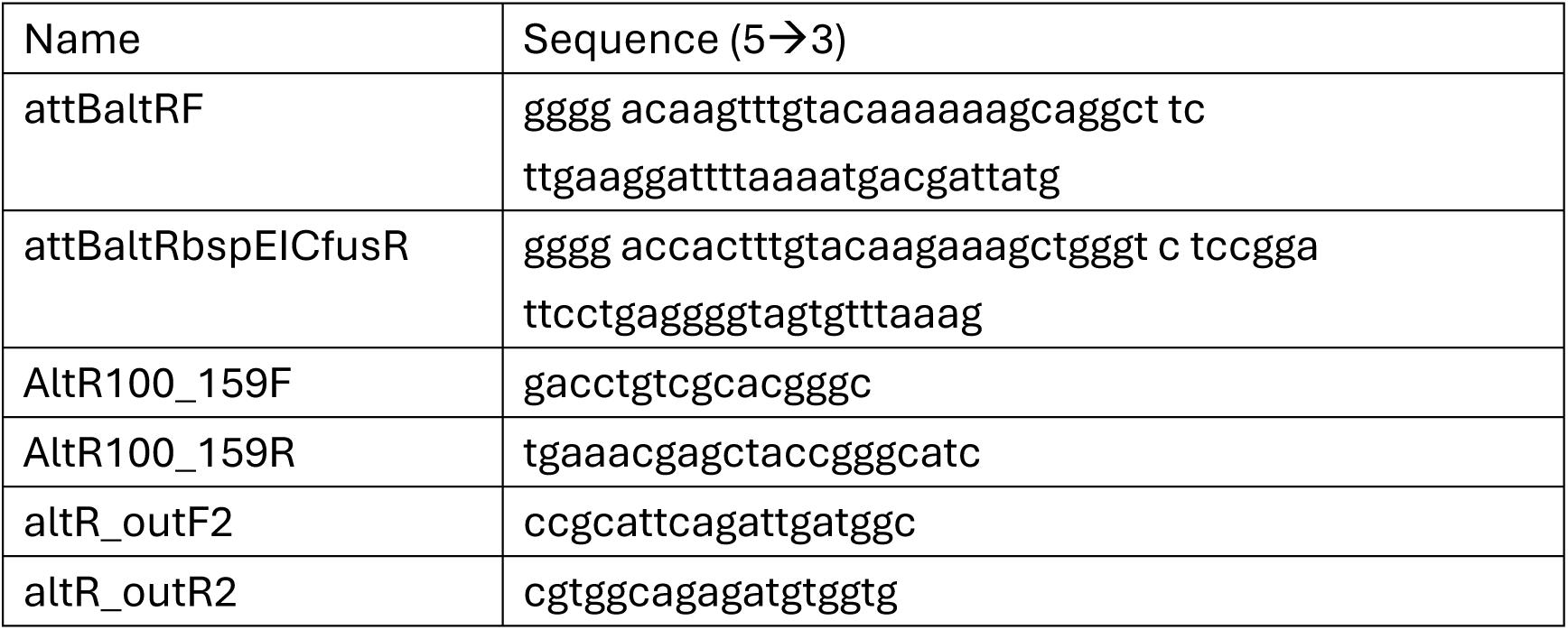
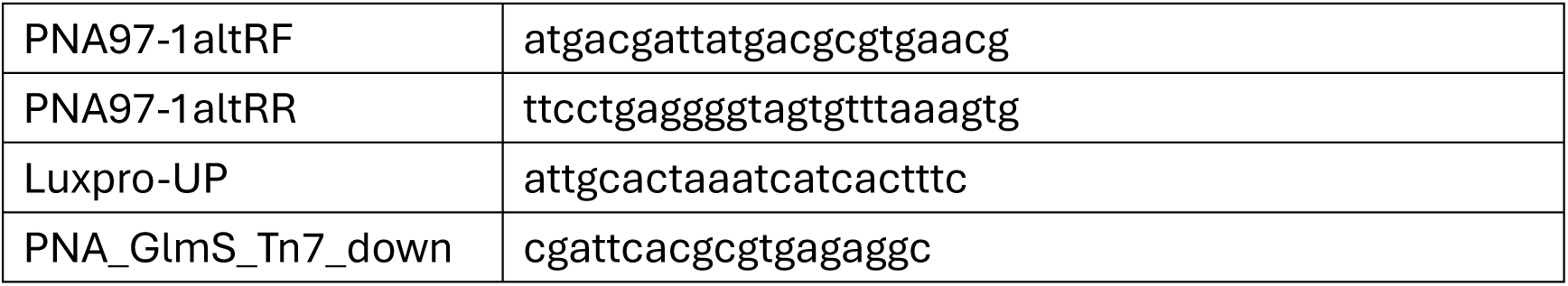
Primers listed in this study for PCR and cloning purposes.

**Table S3.**
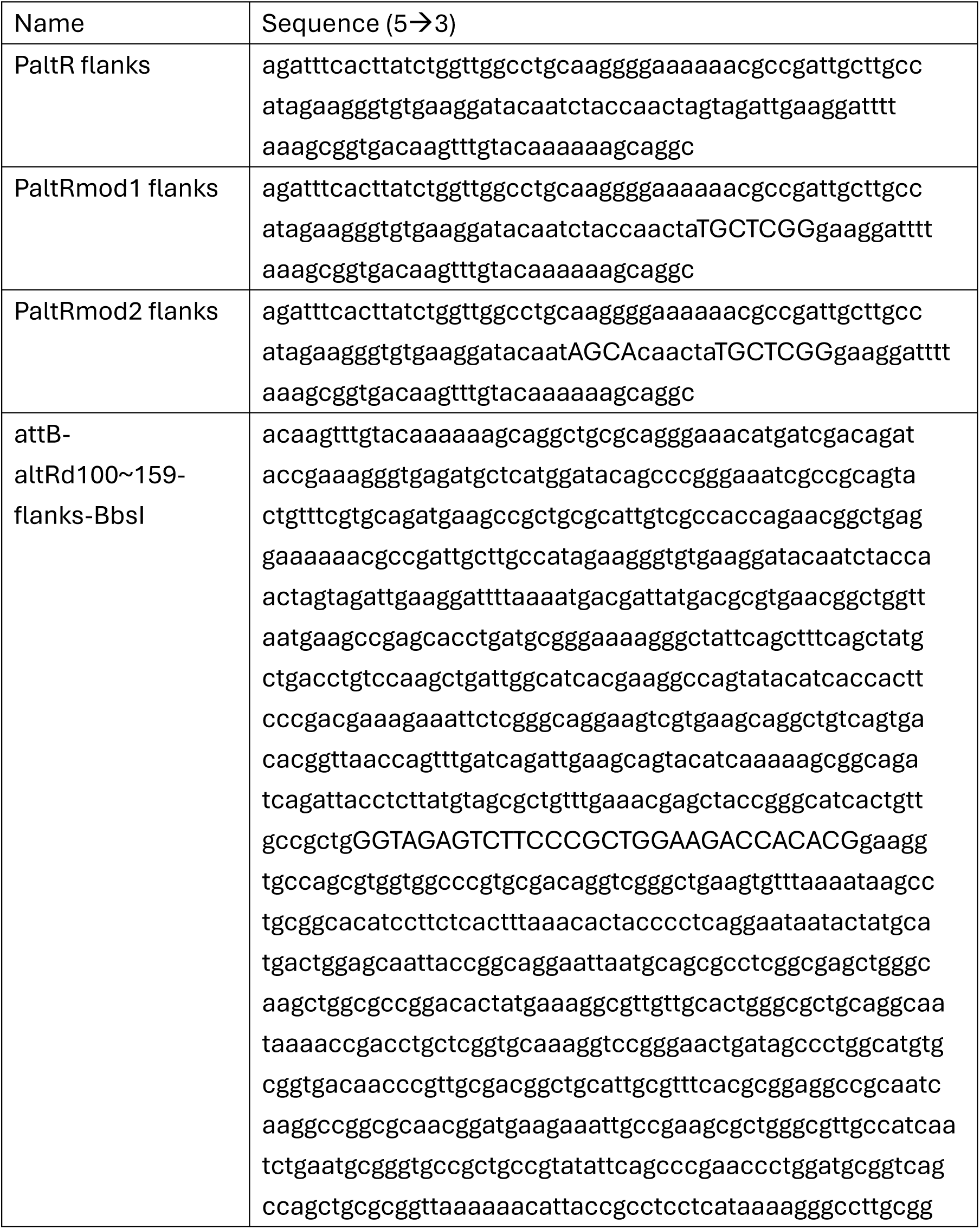
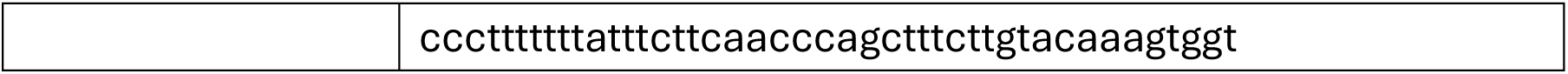
Synthesized sequences that were utilized for Gateway cloning and Gibson assembly in this study.

